# The episodic resurgence of highly pathogenic avian influenza H5 virus

**DOI:** 10.1101/2022.12.18.520670

**Authors:** Ruopeng Xie, Kimberly M. Edwards, Michelle Wille, Xiaoman Wei, Sook-San Wong, Mark Zanin, Rabeh El-Shesheny, Mariette Ducatez, Leo L. M. Poon, Ghazi Kayali, Richard J. Webby, Vijaykrishna Dhanasekaran

**Affiliations:** School of Public Health, LKS Faculty of Medicine, The University of Hong Kong, Hong Kong, China; HKU-Pasteur Research Pole, LKS Faculty of Medicine, The University of Hong Kong, Hong Kong, China; Sydney Institute for Infectious Diseases, School of Life and Environmental Sciences and School of Medical Sciences, The University of Sydney, Sydney, New South Wales, Australia; Department of Microbiology and Immunology, at the Peter Doherty Institute for Infection and Immunity, The University of Melbourne, Melbourne, Victoria, Australia; Centre for Immunology & Infection, Hong Kong Science and Technology Park, New Territories, Hong Kong S.A.R., China; Center of Scientific Excellence for Influenza Viruses, National Research Centre, Egypt; IHAP, Université de Toulouse, Institut national de recherche pour l’agriculture, l’alimentation et l’environnement, Ecole Nationale Vétérinaire de Toulouse, Toulouse, France; Human Link, DMCC, Dubai, United Arab Emirates; Department of Infectious Diseases, St Jude Children’s Research Hospital, Memphis, USA

## Abstract

Highly pathogenic avian influenza (HPAI) H5N1 activity has intensified globally since 2021, replacing the dominant clade 2.3.4.4 H5N8 virus. H5N1 viruses have spread rapidly to four continents, causing increasing reports of mass mortality in wild birds and poultry. The ecological and virological properties required for future mitigation strategies are unclear. Using epidemiological, spatial and genomic approaches, we demonstrate changes in the source of resurgent H5 HPAI and reveal significant shifts in virus ecology and evolution. Outbreak data indicates key resurgent events in 2016/17 and 2020/21 that contributed to the panzootic spread of H5N1 in 2021/22, including an increase in virus diffusion velocity and persistence in wild birds. Genomic analysis reveals that the 2016/17 epizootics originated in Asia, where HPAI H5 reservoirs are documented as persistent. However, in 2020/21, 2.3.4.4b H5N8 viruses emerged in domestic poultry in Africa, featuring several novel mutations altering the HA structure, receptor binding, and antigenicity. The new H5N1 virus emerged from H5N8 through reassortment in wild birds along the Adriatic flyway around the Mediterranean Sea. It was characterized by extensive reassortment with low pathogenic avian influenza in domestic and wild birds as it spread globally. In contrast, earlier outbreaks of H5N8 were caused by a more stable genetic constellation, highlighting dynamic changes in HPAI H5 genomic evolution. These results suggest a shift in the epicenter of HPAI H5 beyond Asia to new regions in Africa, the Middle East, Europe, and North and South America. The persistence of HPAI H5 with resurgence potential in domestic birds indicates that elimination strategies remain a high priority.

---

Influenza A viruses (genus *Alphainfluenzavirus*, family *Orthomyxoviridae*) are negative-sense, single-stranded, segmented RNA viruses categorized into subtypes based on the antigenicity of their two surface proteins. Sixteen of eighteen known hemagglutinin (HA) subtypes and nine of eleven neuraminidase (NA) subtypes are prevalent as low pathogenic avian influenza (LPAI) viruses in wild aquatic birds worldwide.

Highly pathogenic avian influenza (HPAI) viruses evolve from LPAI viruses in poultry by acquiring insertions in the HA cleavage site that facilitates systemic infection (1, 2). Only H5 and H7 subtypes have evolved into HPAI (3), with most outbreaks being contained through culling (stamping out) or die-offs with limited opportunities to reinfect and adapt to wild birds (3). Although an HPAI H5N3 epizootic was reported in wild terns in South Africa as early as 1961 (2), the HPAI H5N1 virus that emerged in China during 1996 (now referred to as the goose/Guangdong (gs/Gd) lineage) (4, 5) was the first documented HPAI lineage to establish sustained transmission in domestic bird networks in Asia. The early evolution of the gs/Gd lineage was characterized by the diversification of the H5 HA gene into as many as ten major phylogenetic clades (6, 7), which through extensive reassortment with LPAI viruses, acquired new combinations of internal genes. Ultimately, clade 2 proved most successful, and its repeated reinfection of wild aquatic birds (8-11) has enabled episodic dissemination from its origins in China to other parts of Asia, Europe, Africa, and most recently, North and South America (5, 12-15).

The scale of wild bird outbreaks due to HPAI H5 has increased outside Asia since 2014 (16-18) and has been driven by the emergence of H5 HA lineage 2.3.4.4 viruses with multiple NA types including H5N2, H5N6 and H5N8 (collectively H5Nx) (**Fig 1a**) (19). Prior to clade 2.3.4.4, HPAI H5 evolution was characterized by a relatively stable internal cassette that mainly derived genes from the avian influenza gene pool in domestic birds (e.g., H9N2) and the linkage of its surface proteins, H5 HA and N1 NA, despite HA diversification. Experimental animal infections showed clade 2.3.4.4 viruses were associated with increased adaptation and reduced virulence in wild ducks compared to earlier H5N1 lineages (20-23). Repeated outbreaks in wild birds from 2016 were due to clade 2.3.4.4.b H5N8 viruses that originated in China (16) and had a relative increase in virulence in ducks (24, 25). Furthermore, a reassortant HPAI H5N1 subtype which evolved from clade 2.3.4.4b viruses, has almost entirely replaced clade 2.3.4.4.b H5N8 viruses (**Fig. 1**). This led to an unprecedented number of infections in wild birds across four continents since November 2021 (26-30), increased poultry infections starting in Europe since early 2020. The associated increase in cases in mammals is predominantly reported from U.S. (31). At least three 2.3.4.4b lineages circulated in 2020 (32). The epidemic source of recent HPAI resurgences, data required to inform future mitigations, underlying genome evolution in wild and domestic populations and its role in virus maintenance and spreading remains unclear. Similarly, the ecological properties that lead to enhanced and sustained transmission in wild birds need to be better understood as they present unpredictable epizootic, zoonotic and pandemic threats in new regions, as do the environmental and evolutionary consequences of poultry vaccination with variable uptake.

**Figure 1.**
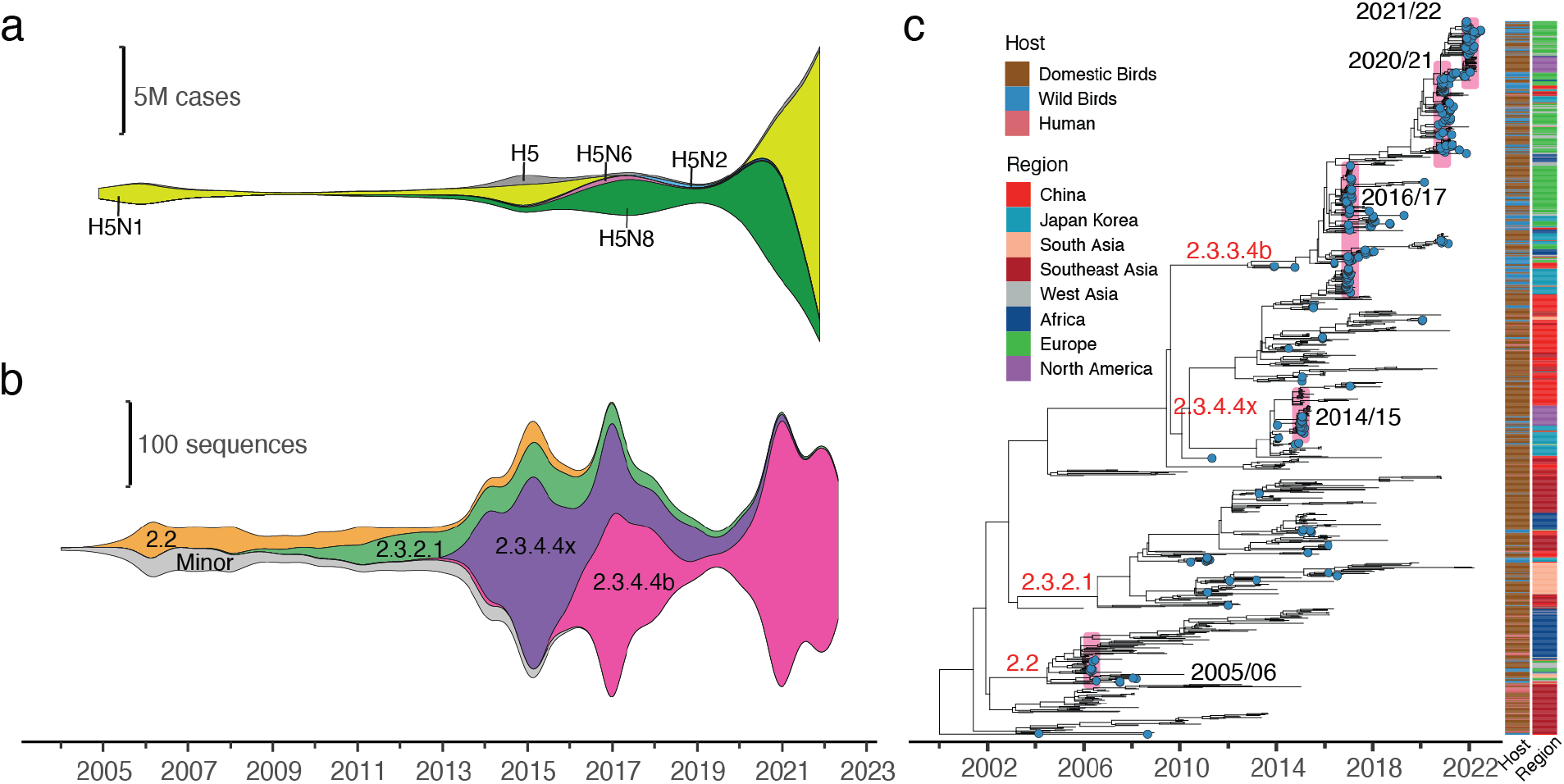
Dynamic changes in HPAI H5 subtypes and clades. **(a) Temporal** changes in HPAI H5Nx subtype prevalence estimated using observation dates of all reported cases submitted to the World Organization for Animal Health (WOAH) from January 2005 to January 2022. **(b)** Temporal changes in HPAI H5 HA clade prevalence estimated using sample collection dates of all AIV sequences submitted to the Global Initiative for Sharing All Influenza Data (GISAID) and NCBI Influenza Virus Resource databases from January 2004 to June 2022. **(c)** Time-scaled maximum likelihood tree of HPAI H5 HA based on 1,000 sequences subsampled from all available AIV sequences (N = 10,643). Four major clades are denoted in red text, and the years of major wild bird resurgence events are highlighted with pink bars and denoted in black text. Tips representing wild bird samples are color-coded in blue. Host and region of isolation are shown as bars.

To address these issues, we examined the changing epidemiology of HPAI H5Nx viruses since 2005 using outbreak data reported to the Food and Agricultural Organization (FAO) and World Organization for Animal Health (WOAH), from which we were able to identify epidemic trends in wild and domestic birds. To investigate how the underlying virus and host demographics are affected by dispersal and genome evolution, we further analyzed over 11,000 whole genomes of HPAI H5 viruses collected globally, spanning these multiple resurgent events, and found substantial shifts in HPAI ecology and evolution.

## Results

### Epidemiology of resurgent HPAI H5N1 outbreaks

The H5 gs/Gd lineage was first detected in 1996, and beginning in 2005, spread substantially through H5N1 clade 2.2 virus outbreaks in poultry and wild birds. Analysis of FAO and WOAH reports since 2005 confirmed a seasonal pattern peaking in winter in both domestic and wild birds in all years (also see below). Reports show four significant HPAI H5 epizootic events in wild birds, characterized by rapid detection of the lineage across multiple continents and increased reports than previous years, reported in the literature due to clade 2.3.4.4 in 2014/15 and clade 2.3.4.4.b in 2016/17, 2020/21, and 2021/22 (**Fig. 2a**) (33-36). H5N8 viruses were the cause of the 2014/15, 2016/17, and 2020/21 outbreaks. While H5N6 was also part of the complex landscape, it played a nominal role in the 2016/17 wild bird resurgence (**Figs. 1a** and **2b**) (37). H5N1 subtype viruses caused the 2021/22 outbreaks (**Figs. 1a** and **2b**), almost completely replacing all other H5 subtypes globally.

**Figure 2.**
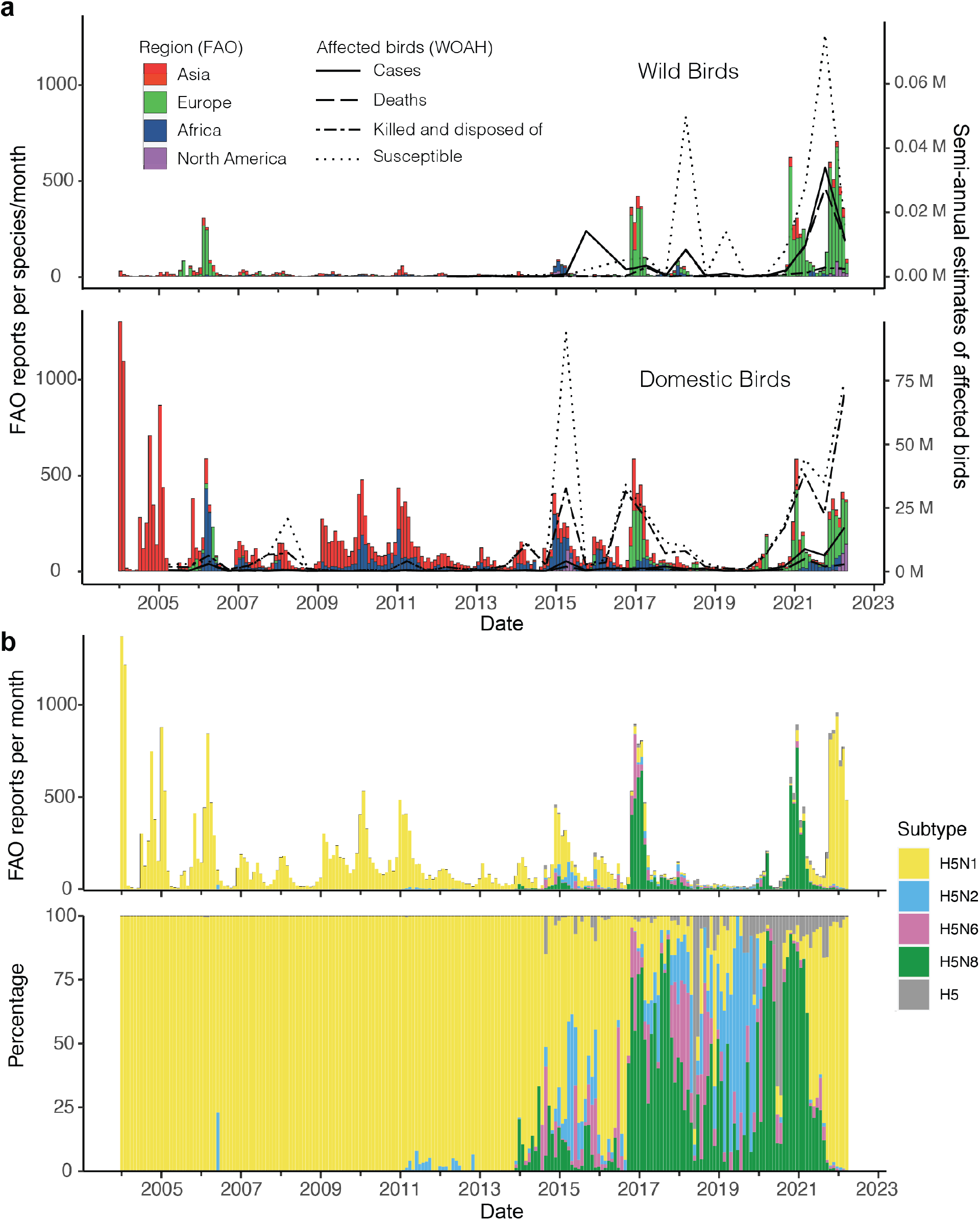
HPAI H5 virus outbreak reports. **(a)** Time series comparing HPAI H5 outbreaks in wild birds (above) and poultry (below) by geographic region reported to the Food and Agriculture Organization of the United Nations (FAO). Semi-annual counts of HPAI H5 affected birds reported to the World Organization for Animal Health (WOAH) are plotted on the right hand y-axis. **(b)** Monthly HPAI H5 outbreaks from January 2004 to May 2022 colored by subtype.

While 2014/15 is a notable resurgence event due to the spread from Asia to Europe and North America (16, 38), resulting in a loss of >50 million poultry in the U.S (39), the number of poultry outbreaks in Europe and wild bird detections globally was low (**Fig. 2a**), possibly due to lower relative virulence and virus fitness. The 2016/17 epidemic in wild birds lasted six months, with over 230 outbreaks reported per month (**Fig. 2a**) (12, 40). The 2017/18 season (September to August) saw fewer reported outbreaks, but a greater number of wild birds were deemed susceptible across multiple regions. Following only sporadic detections from 2018–2020, >200 outbreaks per month were reported in 2020/21, and an unprecedented >400 outbreaks per month were recorded during the 2021/22 season. In addition to the increase in epizootics, an overall increase in the number of wild bird species affected was observed in all regions to varying degrees during the 2021/22 season compared to the previous seasons (Supp fig. 1). The semi-annual number of confirmed HPAI H5 cases in wild birds (predominantly dead birds) peaked at 34,000 during the second half of 2021, although it is important to note that in many instances the number of wild bird cases reported to WOAH/FAO comprises only birds tested and positive for HPAI, which is, therefore, a substantial underestimate (**Fig. 2a**). Increase in the number of domestic bird outbreaks generally corresponded to increases in wild bird outbreaks. Between January– June 2022, over 69 million susceptible domestic birds were culled. Notably, substantial numbers of poultry outbreaks were also recorded in early 2020, prior to the detection of wild bird epizootics, which spiked during April 2020 (**Fig. 2a**).

Long-term outbreak notification data revealed a shift in regional domestic bird outbreak trends. Prior to 2016, outbreaks occurred mainly in Asian and African countries, except for 2005/06 and 2014/15 wherein outbreaks also occurred in Europe and North America (**Fig. 2a**). From 2016–2022, European countries increasingly reported more outbreaks, while reports from Asia and Africa were sporadic, however, the contribution of the declared endemicity of HPAI H5 in these regions to sporadic reporting is unclear. North America reported a substantial number in 2022. A seasonal pattern with increased reports during winter months was observed in Asia since 2005 and in Africa especially during 2014/15, but the cumulative number of reports in these regions remained low. All major wild bird resurgence events in Europe exhibit seasonal patterns, with widespread outbreaks beginning in November (**Fig. 2a** and Supp fig. 2). The seasonality is explained by the arrival of wild migratory birds from their arctic breeding areas (41), with arrival in Europe timed to coincide with the 0°C isotherm (42). This consistent seasonality has led reporting agencies to describe annual events in waves, starting in September; however, cases in Europe have continued through the summer of 2022. The number of wild bird outbreaks in Asia increased during the early months of 2017, 2021, and 2022 when widespread detections in Europe were reported.

### Origins and expansion of resurgent HPAI H5 outbreaks

A comprehensive analysis using all available HPAI H5 genomes shows the 2014/15 and 2016/17 resurgent events originated from independent viral lineages first detected in China (**Figs. 1c** and **3a**). In contrast, all eight genes involved in the 2020/21 outbreaks, which caused a number of epizootics across Eurasia and Africa, originated in 2019 from H5N8 viruses first detected in poultry in Egypt in 2016/17 (**Fig. 3**, Supp figs. 3 and 4 and Supplementary Data 1). Although HPAI H5 surveillance is sparse in poultry networks surrounding Egypt, the continuous detection of ancestral lineages of the 2020/21 resurgent viruses in domestic birds in Egypt strongly suggests the evolution and resurgence of the lineage in the region. The H5N1 viruses responsible for the 2021/22 outbreak, emerged from H5N8 viruses in Europe in 2020 (mean tMRCA of HA gene 17 August 2020; 95% HPD 13 June 2020, 14 October 2020), conforming with epidemiological analysis by European Food Safety Authority which showed the first H5N1 outbreaks were reported as early as October 2020 (43). The N1-NA gene and five internal genes (PB2, PB1, PA, NP, and NS) of H5N1 were newly derived through reassortment from LPAI prevalent in the European wild bird populations since 2019 (Supplementary Data 1). Estimates of tMRCA showed ancestral genes of all three major resurgent lineages (clade 2.3.4.4b H5N8 in 2016/17 and 2020/21 and clade 2.3.4.4b H5N1 in 2021/22) circulated in wild birds for at least a year before causing widespread outbreaks from November onwards.

**Figure 3.**
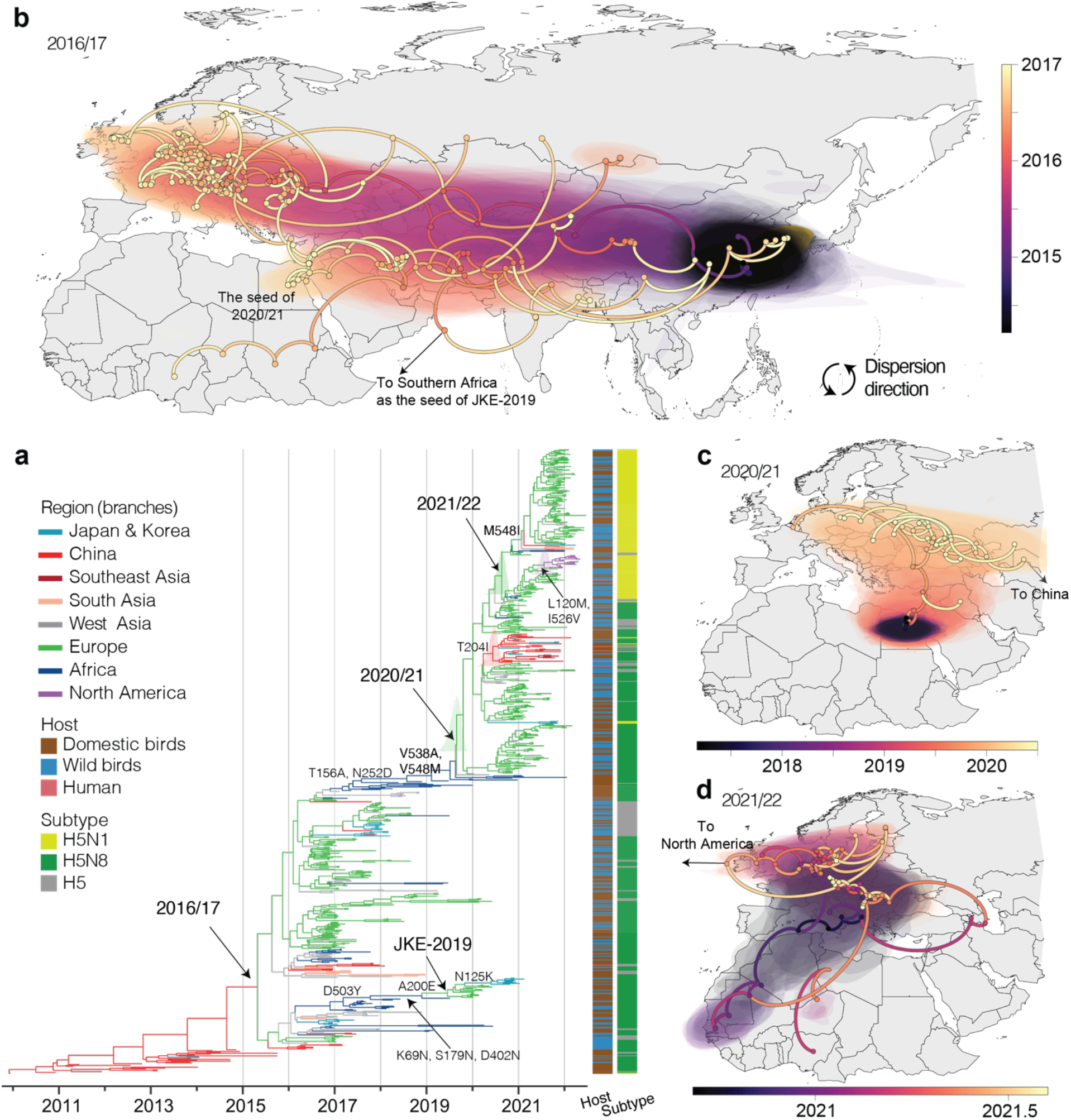
Evolution of clade 2.3.4.4b and early migration patterns of resurgence HPAI. **(a)** Maximum clade credibility tree with branches colored by geographic region. Colors on the right indicate host and virus subtype. The posterior distribution of the time to the most recent common ancestor (tMRCA) is shown as bar charts at specific nodes. **(b)** Continuous phylogeographic reconstruction of the spread of H5N8 from mid-2014 to 2017, **(c)** the early spread of H5N8 from 2017 to mid-2020 before the 2020/21 resurgence, and **(d)** the early spread of H5N1 from mid-2020 to 2021 before the 2021/22 resurgence. Circles represent nodes in the maximum clade credibility phylogeny, colored by the inferred time of occurrence. An interval of 95% higher posterior density is depicted by shaded areas, illustrating the uncertainty of the phylogeographic estimates.

A separate H5N8 clade 2.3.4.4b virus lineage that derived from the 2016/17 resurgence (hereafter termed JKE-2019) was initially reported in domestic poultry in Central and Southeastern Europe during 2019 and early 2020 and later emerged in wild birds in Japan and Korea in 2020/21 (**Figs. 3a-b**). JKE-2019 retained six of eight genes of clade 2.3.4.4b viruses found in West and Southern Africa during 2018–2019, while the PB1 and NP genes were obtained from LPAI viruses through reassortment (17) (Supp fig. 4).

We reconstructed the diffusion of 2.3.4.4b H5N8 viruses using a continuous phylogeographic analysis of HA (**Fig. 3c** and Supp fig. 5) and found that the initial migration from its inferred ancestors in Egyptian domestic poultry was through the Black Sea-Mediterranean flyway towards western Russia and Eastern Europe during 2019, following a pattern described during previous resurgence events (44, 45). During mid-late 2019, an HPAI H5 lineage was introduced in the northern coastal regions of Central Europe, where it circulated locally and reassorted with LPAI until resurgence as HPAI H5N8 in late 2020 and as HPAI H5N1 in late 2021 (**Fig. 3c** and Supp fig. 2). HPAI H5N8 spread rapidly across Eurasia and became established in wild birds and poultry systems during 2020. By October 2020, clade 2.3.4.4b HPAI H5N8 viruses with an additional HA mutation (T204I) migrated east to China, where they acquired an N6-NA, forming an Asian lineage (tMRCA 9 June 2020; 95% HPD 3 April to 11 August), which has been responsible for 41 of the 65 known human cases of H5N6 infection to date (46).

Continuous phylogeography revealed 2.3.4.4b HPAI H5N1 emerged along the Adriatic flyway during mid-2020, with early detections along the Balkan and Apennine peninsulas extending southwest towards the coastal regions in the Northwest and West Africa, as well as northeast to form two separate lineages in northern Central and Eastern Europe, respectively (**Figs. 3b** and **3d**). A lineage introduced into the northern coastal regions of Central Europe in late 2020 (mean tMRCA of HA gene 27 November 2020; 95% HPD 30 September 2020, 24 January 2021) was maintained across northern Europe and seeded North American outbreaks with two additional HA mutations (L120M and I526V) in mid-2021 (mean tMRCA of HA gene 24 July 2021; 95%HPD 27 May, 19 September). Another lineage circulating in Europe acquired an additional HA M548I substitution before causing outbreaks across Eurasia during 2021/22, suggesting the potential importance of the site for adaptation to rapid dispersal among wild birds.

### Contrasting phylogeography and phylodynamics of clade 2.3.4.4b viruses

The emergence of multiple lineages from regions outside Asia signifies a significant shift of the gs/Gd HPAI H5 epicenter. To infer changes in spatial dispersal and cross-species dynamics of gs/Gd HPAI H5 among wild and domestic bird populations, we applied an asymmetric discrete-trait model with Bayesian Stochastic Search Variable Selection (BSSVS) to reconstruct diffusion of major gs/Gd HPAI H5 clades among eight geographic regions (Africa, China, Europe, Japan and Korea, North America, South Asia, Southeast Asia, and West Asia) and different hosts (domestic and wild birds). In contrast to the predominant HPAI H5 clades that circulated in previous years (2.2, 2.3.2.1 and other 2.3.4.4 subclades including a,c–f, collectively 2.3.4.4x), which primarily involved regional circulation in domestic bird populations with brief periods of rapid dispersal via wild birds (**Fig. 4**), clade 2.3.4.4b viruses are characterized by a considerably longer mean duration of persistence in wild birds (51.8% Markov rewards, denoting time spent in the host) than domestic birds, with 76.4% of Markov jumps (denoting host transitions) going from wild birds to domestic birds (**Fig. 4** and Supp fig. 6), and an epicenter shift from Asia to Europe. Specifically, significant source populations of clade 2.3.4.4b were in Europe (43 Markov jumps, 44.5%), West Asia (22 Markov jumps, 23.3%), and China (11 Markov jumps, 12%) (Supp fig. 6). Lineages spent the most time in Europe (51.7% Markov rewards), followed by China (16.7% Markov rewards) and West Asia (10.9% Markov rewards), whereas Africa (13.6% Markov rewards) acted as a sink (Supp fig. 6), as recently shown (47). However, clade 2.3.4.4b has persisted in Europe and Africa for around two years, compared to roughly one year in other regions. Furthermore, while domestic birds in China occupied the phylogenetic trunk until 2015 and were the probable source of the 2016/17 wild bird outbreaks, Europe formed the primary phylogenetic trunk since 2016 and transmission was primarily among wild birds, except around 2019 when over 50% of the phylogenetic tree trunk was occupied by lineages circulating in domestic birds in Africa which contributed to resurgent epizootics since 2020 (**Fig. 4**). HA phylogenies show several sub-lineages derived from the 2016/17 wild bird resurgence in Europe spread to Africa and several of these were maintained in domestic and wild bird populations detected in multiple countries including South Africa, Nigeria, however, five of nine lineages detected in 2019/20 were from Egypt (**Fig. 3a**, https://nextstrain.org/community/vjlab/episodic-h5/H5). Sustained transmission of 2016/17-like viruses was also detected in Iran, Denmark and Bulgaria until 2019, whereas non-2.3.4.4b clades remained predominant in Asia until 2020.

**Figure 4.**
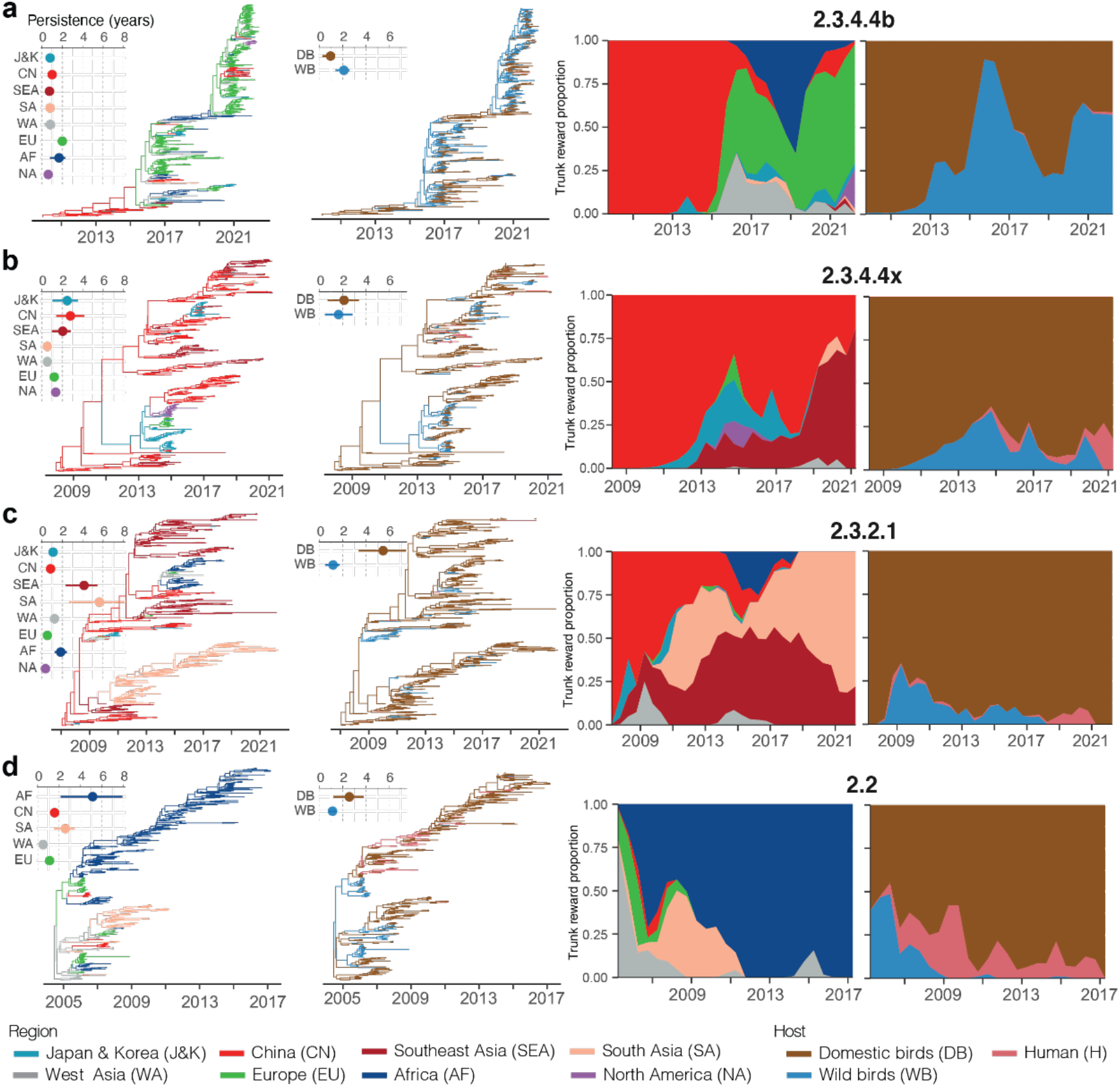
The contrasting geographic and host transmission patterns of major HPAI H5 clades including 2.3.4.4b (a), 2.3.4.4x (b), 2.3.2.1 (c) and 2.2 (d). From left to right, the figures represent Maximum clade credibility trees for each clade colored by eight geographic regions, including Japan and Korea (J&K), China (CN), Southeast Asia (SEA), South Asia (SA), West Asia (WA), Europe (EU), Africa (AF) and North America (NA) in the first column and hosts including domestic birds (DB), wild birds (WB) and human (H) in the second column. To the top left of each tree is an inset that measures the duration of region/host-specific persistence in years. Each circle represents the mean persistence of the sampled viruses, and each line represents the inter-quartile range of persistence. The third and fourth columns show proportional ancestral region and host states on the phylogenetic tree trunk over time.

The longest migration routes from China to the United States via Europe and frequent transmission via wild birds between Asia and Europe observed for 2.3.4.4b were rare for other clades (Supp fig. 5). Clade 2.3.4.4x was frequently disseminated from China to Southeast Asia, Japan and Korea (over five Markov jumps), and once to North America, whereas clade 2.3.2.1 migration between China and Southeast Asia over five Markov jumps, and to a lesser extent, to Japan and Korea and Africa via West Asia (Supp figs. 5 and 6). Following the early dispersal of clade 2.2 to Europe and Africa (48), dispersal between West Asia to Africa via Europe was mainly through domestic bird networks, while wild birds were primarily responsible for transmission between West Asia and Europe. Our analysis suggests the majority of regional migration of clades 2.2, 2.3.2.1 and 2.3.4.4.x were likely via domestic birds and associated human activity (Supp fig. 5); however, short-range spread via unsampled wild birds cannot be excluded.

To better understand the spatial epidemiology of resurgent HPAI H5 clades, we estimated the wavefront distance and velocity (49) from a joint phylodynamic analysis that incorporated continuous spatial diffusion and discrete avian hosts (domestic/wild) (50) (**Fig. 5** and Supp figs. 5 and 7). These estimates showed that the diffusion velocity and coefficient in wild birds are faster than that of domestic birds, as expected for all major gs/Gd clades. However, in contrast to previous studies that showed steady range expansion of pre 2.3.4.4 HPAI H5 due to domestic birds as measured by the wavefront distance (11), we find wild birds contributed to a greater geographic expansion of both clade 2.3.4.4 lineages (3,630 km (95% HPD: 1381 – 6977) for 2.3.4.4b and 4,517 km (95% HPD: 0 – 8392) for 2.3.4.4x) compared to clades 2.2 (2199 km, 95% HPD: 2 – 5097) and 2.3.2.1 (526 km, 95% HPD: 0 – 1364). Furthermore, wild and domestic birds contributed almost equal wavefront distance (∼5,000 km) since 2.3.4.4.b resurgence in wild birds since late 2019 (termed panzootic-2020 in **Fig. 5**). However, in 2020, the diffusion velocity of 2.3.4.4b from wild birds was nearly three times (5,474 km/year; 95% HPD: 3,535 – 7,877) that of domestic birds, after which the diffusion velocities in wild and domestic birds were similar (3,000 – 4,000 km/year).

**Figure 5.**
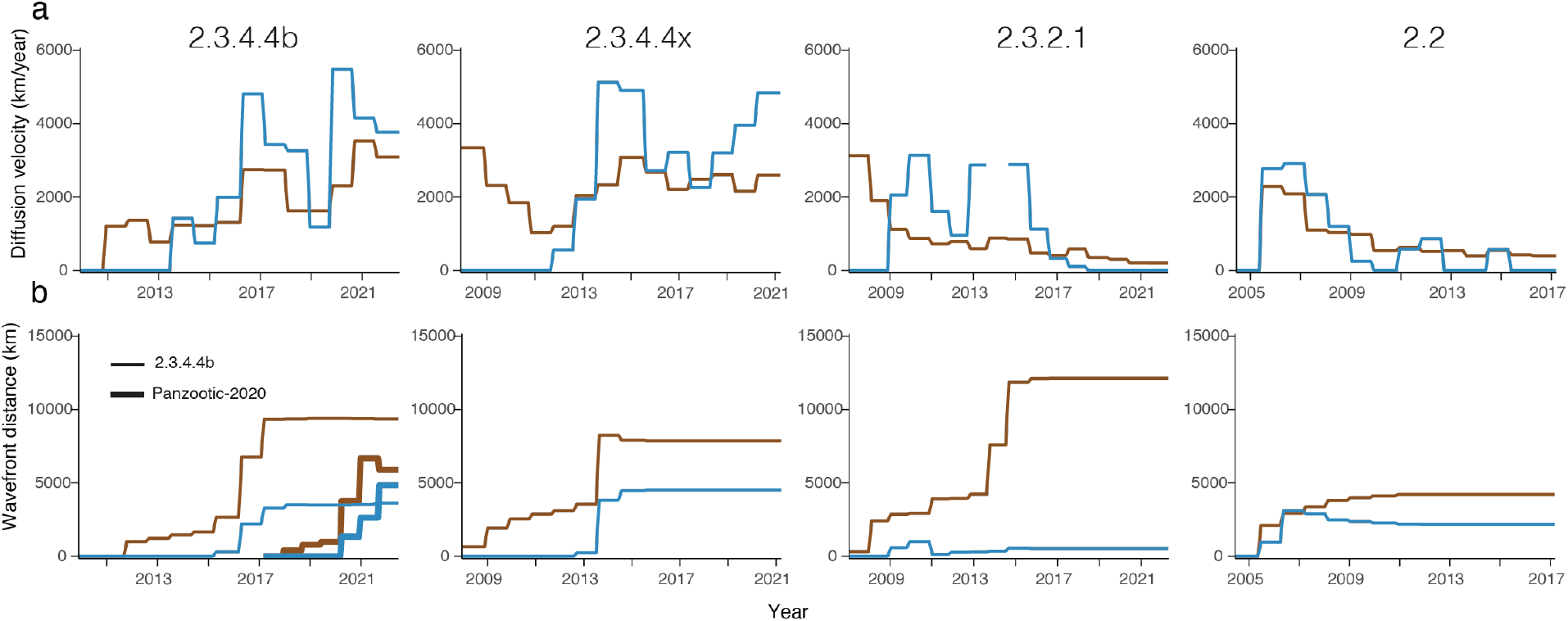
The contrasting spatial epidemiology among HPAI H5 clades 2.3.4.4b, 2.3.4.4x, 2.3.2.1 and 2.2. **(a)** Viral Diffusion velocity (km/year) of domestic (brown) and wild birds (blue) over time for each clade. The diffusion velocity of wild birds in clade 2.3.2.1 during 2014 is not shown in the plot due to abnormal estimation value, possibly caused by insufficient sampling. **(b)** Viral wavefront distance (km) of wild and domestic birds over time for each clade. A recalculation of the wavefront distance in the panzootic-2020 clade (including 2020/21 and 2021/22 resurgences, shown in a thick line) was performed, in which Egypt was regarded as the epidemic’s origin. Confidence intervals are shown in Supp fig. 7.

### Contrasting patterns of reassortment in wild birds and antigenic evolution in poultry

The early adaptation of HPAI H5N1 to domestic poultry between 1996–2004 was characterized by extensive reassortment with the LPAI gene pool in domestic poultry, such as H9N2 viruses adapted to poultry, with few gene incursions from the LPAI gene pool circulating in wild birds across Eurasia (51). In contrast, the emergence and circulation of clade 2.3.4.4 was characterized by increased reassortment (Supp fig. 8), deriving multiple NA subtypes and internal genes from the greater gene pool diversity prevalent in wild birds across Eurasia (16). Strikingly, the vast majority of resurgent HPAI H5N8 viruses collected in 2020/21 maintained a stable genotype with all genes originating from poultry in Egypt, except a lineage in China in which N1 and N6 NA genes were more frequent. However, prior to the wild bird resurgence, multiple HA mutations occurred in domestic birds in Egypt between 2017–2020, including T156A and N252D mutations in antigenic regions; the former has been shown to enhance the overall binding of HA to sialic acid receptors (52), as well as V538A and V548M mutations in the conserved HA stalk region before emergence (**Fig. 3a**). In contrast, the novel 2.3.4.4b HPAI H5N1 that emerged from 2.3.4.4b HPAI H5N8 along the Adriatic flyway repeatedly acquired internal genes through reassortment with LPAI (**Fig. 6**). Novel 2.3.4.4b HPAI H5N1 reassortants increasingly emerged during 2020/21 and 2021/22, and several genotypes generated in 2020/21 persisted to seed outbreaks during 2021/22 (**Fig. 6**). These results suggest that a predominantly clonal population of HPAI H5N8 caused the 2020/21 outbreaks. In contrast, the novel H5N1 that emerged in wild birds attained relaxed constraints for reassortment with LPAI in comparison to ancestral HPAI H5N8.

**Figure 6.**
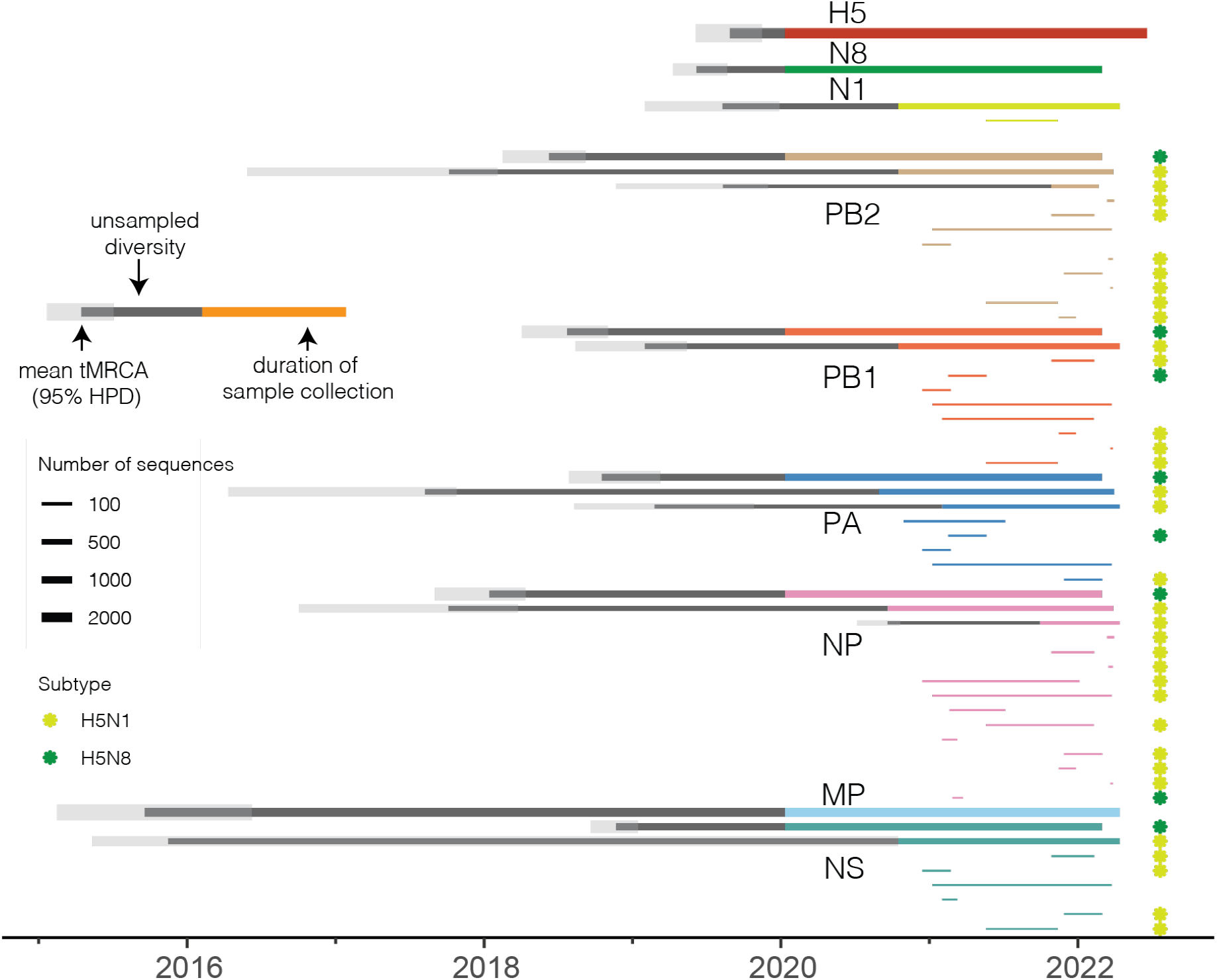
Persistence of HPAI gene segments in resurgent 2020/21 and 2021/22 viruses. Line thickness indicates the number of sequences in a monophyletic clade (with a minimum of five sequences). The internal genes of H5N1 and H5N8 were labeled as yellow and green snowflakes separately after each line. The tMRCA is shown for monophyletic clades with over 100 sequences wherein BEAST v1.10.4 (53) was used for the HA gene and TreeTime (54) for the remaining genes.

## Discussion

A resurgent clade 2.3.4.4b virus caused large-scale outbreaks in wild and domestic birds since late 2020, with increasing severity and panzootic spread since the 2021/22 season. For the first time, human infections have been detected in areas where the gs/Gd H5 virus is not endemic in domestic birds, such as the United Kingdom, the United States, and Russia. At the same time, 2.3.4.4b HPAI H5N1 continues to spread to new countries, with the most recent reports emerging from Peru, Ecuador and Venezuela in November 2022. By analyzing sequence data and outbreak records, we identified the origins of ongoing epizootics as being 2.3.4.4b HPAI H5N8 viruses detected in poultry in Egypt following a major wild bird resurgence event in 2016/17 that originated in China. The acquisition of HA mutations in domestic birds in Egypt, where vaccination is used, that may facilitate endemicity in poultry and allow escape of potential immunity in the wild bird population reinforces evidence that novel HPAI H5Nx variants with significant resurgence potential are repeatedly generated in poultry. Although the viruses show greater persistence in wild birds, our results suggest that continual elimination of HPAI H5Nx in poultry remains a high priority and provides evidence that new HPAI viruses are now emerging outside regions previously thought to drive virus emergence. These results reveal major changes in HPAI H5 ecology and evolution.

Our analysis of WOAH/FAO notification data and genome sequences suggests that wild bird populations in northern Europe may have harbored poultry-derived 2.3.4.4b HPAI H5N8 and H5N1 viruses for over a year prior to their resurgence in the early winter of 2020 and 2021. Previous work has shown that local amplification of HPAI H5 occurred in the Netherlands during the 2016/17 resurgence (55). The seasonality and epidemiology of influenza viruses is closely linked to wild bird ecology. For example, in instances where wild birds, particularly wild ducks, have experienced successful breeding due to favorable climatic conditions, there is substantial juvenile recruitment into the population such that >30% of the population may be susceptible (immunologically naïve) prior to aggregation and migration (56, 57). There is also evidence that many migratory bird species have been affected by recent climatic variations (58, 59). Additionally, an increase in susceptible individuals will also occur in cases wherein the pathogen removes exposed individuals from the population through processes such as migratory culling (60, 61). Potential increases in wild bird population sizes linked to high levels/rates of juvenile recruitment in these regions may thus enable such sustained transmission. However, the full extent of wild bird-domestic poultry viral exchange is not fully understood due to underreporting of events in many countries where avian influenza surveillance in various bird populations is not conducted. This is true for most countries, especially low- and middle-income countries. The events recorded in Egypt may be occurring in other locations where poultry-wild bird interfaces exist. Notably, in contrast to earlier work that showed Africa mainly acted as a sink for HPAI H5 viruses (47), our findings show a dynamic shift in the ecological source of clade 2.3.4.4.b from Asia to Africa in recent years, with at least one lineage detected outside Egypt spreading across multiple continents. This highlights the importance of reinforcing surveillance efforts in Africa.

Our analysis showed dynamic changes in AIV evolution in domestic poultry and wild birds. The 2.3.4.4b H5N8 virus that caused the 2020/21 outbreaks across Eurasia and Africa was constrained for reassortment in wild birds. In contrast, the novel panzootic 2.3.4.4b H5N1 virus is associated with extensive reassortment with LPAI viruses, indicating a greater potential for enhancing transmission and maintenance in wild birds. Generally, while mammalian influenza viruses support a limited number of stable genome constellations, LPAI viruses in wild birds, particularly ducks, are more frequently reshuffled by reassortment to form transient genome constellations with no clear pattern of gene segment association (62). Some levels of stable circulation of genotypes have been noted in gulls. In contrast, poultry-adapted LPAI viruses show the genetic linkage of the HA and NA segments (e.g., H9N2, H7N9), but exchange the internal gene segments (63). The increased propensity of reassortment of recent HPAI HA with other influenza viruses prevalent across domestic poultry, such as the recently observed reassortment between clade 2.3.4.4. and H9N2 in Burkina Faso (64), is a concern.

Sustained epizootics in wild aquatic birds with repeated spillover and spillback from domestic birds increase the zoonotic and pandemic risks (32). In response to the uncontrollable spread of the 2021/22 wave, poultry vaccination is increasingly being considered in Europe and North America to reduce the likelihood of endemicity. Several countries in Asia and Africa currently use vaccination to mitigate HPAI in poultry with variable impacts (65), in contrast to, Hong Kong, parts of China and Southeast Asia which use vaccination in combination with other methods to eliminate AIV (66, 67). A key concern is the role of poultry vaccination in driving endemicity and the evolution of antigenically diverse HPAI H5 lineages (68), as observed for two HPAI H5N8 lineages in Africa that have caused epidemics since 2020. The proximity of poultry networks to major flyways, such as those in Northern Africa, the Middle East and Eastern Europe, with increased use of diverse vaccination, is also concerning. To address these issues, it is necessary to enhance global surveillance and develop a multi-faceted mitigation strategy tied to this surveillance. There is also a need to improve the potency and cross reactivity of existing H5 vaccines for poultry.

## Methods

### Data source and preparation

HPAI H5 genomes with at least HA gene and sample information, including sample collection date, location, and host species, were obtained from the Global Initiative on Sharing All Influenza Data (GISAID) (https://platform.epicov.org/) and NCBI Influenza Virus Resource (https://www.ncbi.nlm.nih.gov/genomes/FLU/) databases on 11 July 2022. After removing duplicate isolates, laboratory-derived and mixed-subtype isolates, sequences with less than 85% gene coverage (69), and sequences with incomplete collection dates, H5 clades based on the WHO gs/Gd H5N1 nomenclature system (7) were determined using LABEL v0.6.3 (70). Clade 2.3.4.4 was further assigned into subclades 2.3.4.4a–h based on phylogenetic relationships to WHO H5 candidate vaccine viruses with known clade assignment estimated using a maximum likelihood phylogenetic tree generated using the Jukes-Cantor nucleotide substitution model in FastTree v2.1.1.

Avian hosts were classified as domestic and wild birds using strain names, associated metadata, and original publications (https://github.com/vjlab/episodic-h5). Location coordinates were used to analyze diffusion in continuous space and were accurate, at least to the province/state level for China, Russia, and the United States. For discrete analysis and visualization, locations were classified into countries and regions according to the country-region list in NextStrain (71).

To mitigate sampling bias for phylogeographic analyses, curated datasets were subsampled by either epidemiological information (including country, host, and sampling date) or phylogenetic relationship. HPAI H5 HA datasets were subsampled randomly with at most two sequences for clade 2.3.4.4b, five sequences for clade 2.3.4.4x, three sequences for clade 2.3.2.1 and six sequences for clade 2.2 per country per host per season of each year (season 1, January-March; season 2, April-June; season 3, July-September and season 4, October-December). The final subsampled HA datasets contained 715, 617, 579, and 563 sequences for the four clades.

The R package ‘ggstream’ v.0.1 was used to map temporal changes in the sampling of HPAI H5 clades, and ‘rworldmap’ v.1.3 was used to plot world maps.

### Phylogenetic analyses

Gene sequences were aligned using MAFFT v7.490 (72) and trimmed using trimAL v1.4 (73) with a 50% gap threshold followed by manual trimming to the open-reading frame. Maximum-likelihood (ML) trees were generated using IQ-TREE v2.1.4 (74) with the best-fit nucleotide substitution model. The ML phylogenies were used to check each dataset for molecular clock outliers (sequences that have disproportionately too much or too little root-to-tip divergence for its sampling time) using TempEst v1.5.3 (75). The time-scaled ML trees in this study were performed using TreeTime (54). HA phylogeny showing sampling location (country/region), host and clade details of all sequences are available as NextStrain (71) builds at https://nextstrain.org/community/vjlab/episodic-h5/H5.

### Reassortment analyses

To compare reassortment events that occurred among major clades, including clades 2.3.4.4b, 2.3.4.4x, 2.3.21 and 2.2, we first constructed ML trees for each gene using IQ-TREE v.2.1.4. At most 200 isolates with eight segments were subsampled using the Phylogenetic Diversity Analyzer (PDA) tool v1.0.3 (www.cibiv.at/software/pda) (76) for each major clade and the minor clade, finally resulting in 909 sequences for each HA, PB2, PB1, PA, NP, MP and NS dataset, and 897 sequences for NA dataset, which excluded minor subtypes of NA genes (N3, N4, N5 and N9). Baltic package (https://github.com/evogytis/baltic) was used to visualize the incongruence between phylogenetic trees of eight genes. The HA tree was rooted using clade 0, and the remaining genes were mid-point rooted. Isolates were colored according to the HA clade.

Furthermore, the timeline of reassortment of 2020/21 and 2021/22 viruses was inferred using their divergence times summarized in Supplementary Data 1 and **Fig. 6**.

### HPAI H5 poultry and wild bird outbreaks

All reported and confirmed infections of HPAI H5 viruses in humans were obtained from WHO. Confirmed detections/outbreaks in domestic and wild birds globally were obtained from World Animal Health Information System (WAHIS), World Organisation for Animal Health (https://wahis.woah.org/) and EMPRESS-i+ Global Animal Disease Information System, Food and Agriculture Organisation (https://empres-i.apps.fao.org/).

### Bayesian Evolutionary Inference

Divergence times and evolutionary rates were estimated using an uncorrelated relaxed clock model under a Bayesian framework using Markov chain Monte Carlo (MCMC) sampling in BEAST v1.10.4 package (53) and the BEAGLE high-performance library (77). A flexible Gaussian Markov Random Field Skyride coalescent model and a General Time Reversible nucleotide substitution model were used with a gamma distribution of substitution rates. At least two independent MCMC chains with 100 million states were performed for each lineage, sampling every 20,000 and discarding 10% as burn-in. Runs were combined to ensure an effective sample size (ESS) of over 200 was achieved in Tracer v1.7.1. A subset of 500 trees randomly selected from the posterior distribution was used to generate an empirical tree distribution used in the subsequent phylogeographic analysis, an approach that reduced computational time and burden (11, 50).

### Discrete phylogeography

To reconstruct spatial diffusion among a set of eight geographic regions (Africa, China, Europe, Japan and Korea, North America, South Asia, Southeast Asia, and West Asia) and different hosts (domestic birds, wild birds, and humans), we conducted asymmetric discrete-trait phylogeographic analyses with Bayesian Stochastic Search Variable Selection (BSSVS) in BEAST v1.10.4 (53). SpreadD3 v0.9.6 (78) was used to estimate Bayes factors (BF) to determine how statistically significant specific findings are (‘definitive’ (BF > 100) or ‘sufficient’ (100 > BF > 3)). To count all the transitions between states and the time spent in the states between two transitions, we also applied the continuous-time Markov chains (CTMCs) model (79) to complete the Markov jump history over time. We combined three independent chains with 5 million MCMC steps for each lineage and sampled every 10,000 states. The first 10% of each run was discarded as burn-in, resulting in 1350 posterior trees with estimates of the ancestral region and host for each internal node. The trunk region/host through time was measured from these posterior phylogenies using PACT v0.9.5 (https://github.com/trvrb/PACT), where the trunk consists of all branches ancestral to a virus that was sampled within one year of the most recent sample (80).

### Phylodynamics incorporating geography and host

As a complementary to the discrete spatial diffusion analysis to reconstruct a more detailed geographic history, we estimated the HPAIV H5 diffusion in continuous space (latitude and longitude of country level) using a Cauchy relaxed random walk (RRW) model (81) with 0.001 jitter window size. Moreover, following the Bayesian method proposed by Trovão et al. (50), a continuous spatial diffusion process and a discrete host transmission process was incorporated in a single Bayesian analysis to quantify host-specific diffusion rates and geographic expansion (wavefront distance) for each lineage (https://github.com/vjlab/episodic-h5). At least two independent MCC chains were performed, sampled every 10,000, and discarded 10% as burn-in to ensure ESS >200 for each parameter. The continuous phylogeographic analysis was visualized using the R package ‘seraphim’ (82) and codes from McCrone et al. (83) and Dudas et al. (84) using the python library ‘matplotlib’.

## Supplementary materials

**Supplementary Figure 1** | Comparison of affected wild bird species in H5Nx outbreaks between 2016-2017 and 2021-2022.

**Supplementary Figure 2** | The spatial distribution of FAO outbreaks in Europe during 2020-2022. Dots are colored by subtypes.

**Supplementary Figure 3** | Temporal changes in HPAI H5 lineage predominance. (a) The number of HA sequences colored by lineage since 2020. (b) Proportional lineage distribution by month inferred from (a).

**Supplementary Figure 4** | Evolutionary relationships of Panzootic-2020 (including 2020/21 and 2021/22 resurgence) and JKE-2019 lineage. Maximum likelihood tree of Panzootic-2020 (a) and JKE-2019 (b) for eight segments. Samples collected in Africa are highlighted in red.

**Supplementary Figure 5** | Dynamics of HPAI H5 transmission lineages in clades 2.3.4.4.b, 2.3.4.4.x, 2.3.2.1 and 2.2. Virus lineage movements were inferred by continuous phylogeographic analysis for each clade.

**Supplementary Figure 6** | The contrasting geographic and host transmission patterns among HPAI H5 2.3.4.4.b, 2.3.4.4.x, 2.3.2.1 and 2.2 clades inferred from discrete phylogeography. From left to right, the figures represent regional Markov jumps, regional Markov rewards, host Markov jumps and host Markov rewards.

**Supplementary Figure 7** | The contrasting diffusion coefficient (km2/day) among HPAI H5 clades 2.3.4.4b, 2.3.4.4x, 2.3.2.1 and 2.2. The diffusion coefficient of wild birds in clade 2.3.2.1 during 2014 did not show in the plot due to abnormal estimation value possibly caused by insufficient sampling.

**Supplementary Figure 8** | Tanglegram of HPAI H5 virus reassortment. Colored lines connect each virus across all eight genes, showing incongruence between and within major clades.

**Dataset 1** | Summary of HPAI H5 monophyletic clade of eight segments.

**Dataset 2** | Acknowledgements to sequences obtained from GISAID and NCBI (assessed on 11 July 2022).

## Acknowledgments

The computations were performed using research facilities offered by Information Technology Services, the University of Hong Kong. We gratefully acknowledge the staff from the originating laboratories responsible for obtaining the specimens and the submitting laboratories where the genome data were generated and shared via GISAID (Supplementary Data 2). The funding bodies had no role in the design of the study and collection, analysis, and interpretation of data and writing of the manuscript.

## Author contributions

V.D. designed the research. R.X., and X.W. curated the genome datasets. K.M.E. curated the outbreak reports. R.X., K.M.E., X.W., and V.D. performed analysis and designed the Figures. R.X., K.M.E., M.W., and V.D. wrote the manuscript with input from S-S.W., M.Z., R.E-S., M.D., L.L.M., G. K., and R.W. All authors discussed and approved the manuscript.

## Data and materials availability

All anonymized data, code, and analysis files are available in the GitHub repository (https://github.com/vjlab/episodic-h5).

