## Supplementary figures and images for "The episodic resurgence of highly pathogenic avian influenza H5 virus"

### Supplementary Figure 1

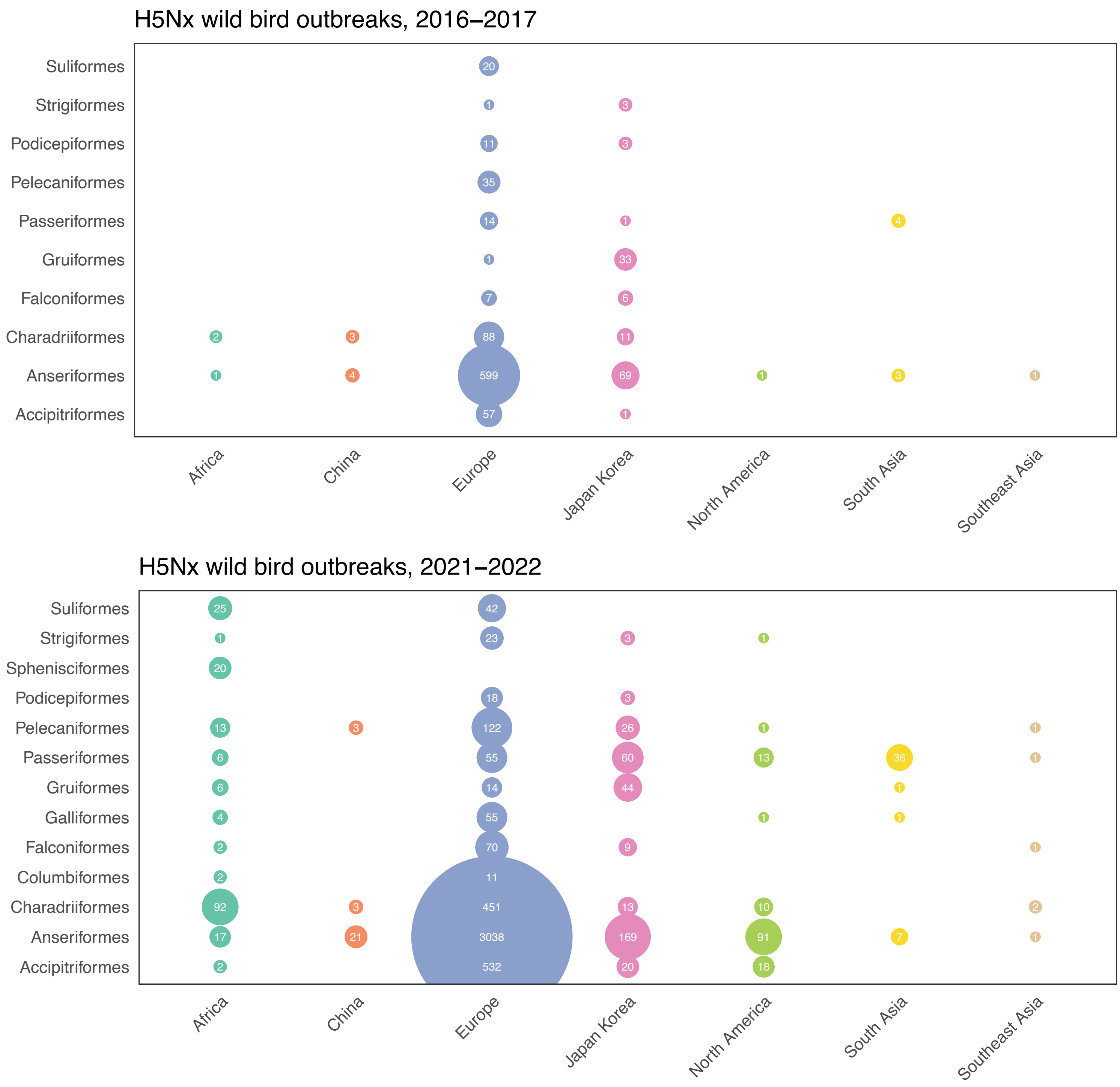


**Supplementary fig. 1.** Comparison of affected wild bird species in H5Nx outbreaks between 2016-2017 and 2021-2022.

### Supplementary Figure 2

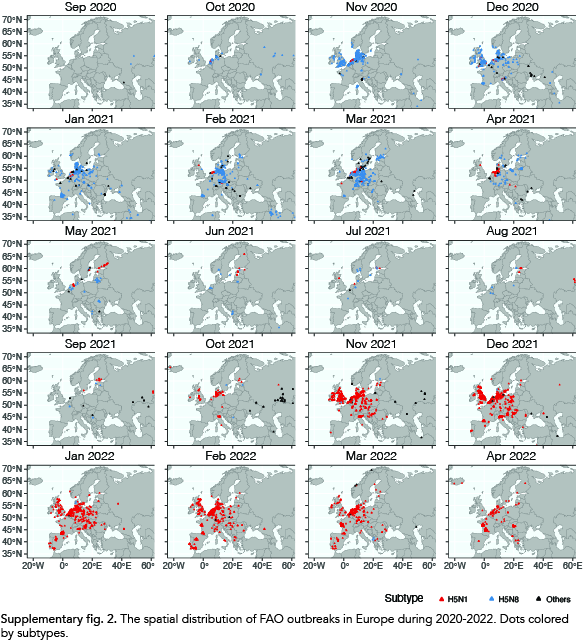
