## Supplementary Figure 3 for "The episodic resurgence of highly pathogenic avian influenza H5 virus"

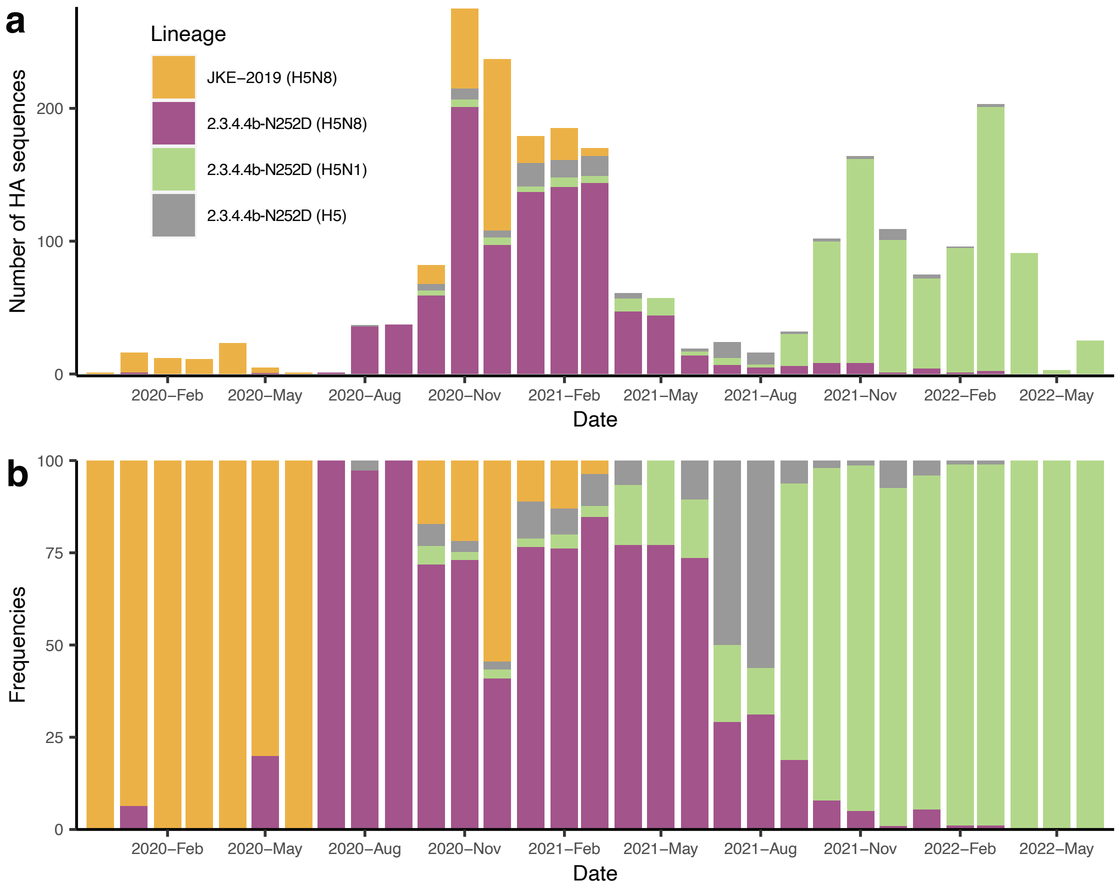


**Supplementary fig. 3.** Temporal changes in HPAI H5 lineage predominance. (a) The number of HA sequences colored by lineage since 2020. (b) Proportional lineage distribution by month inferred from (a).
