## Supplementary Figure 4 for "The episodic resurgence of highly pathogenic avian influenza H5 virus"

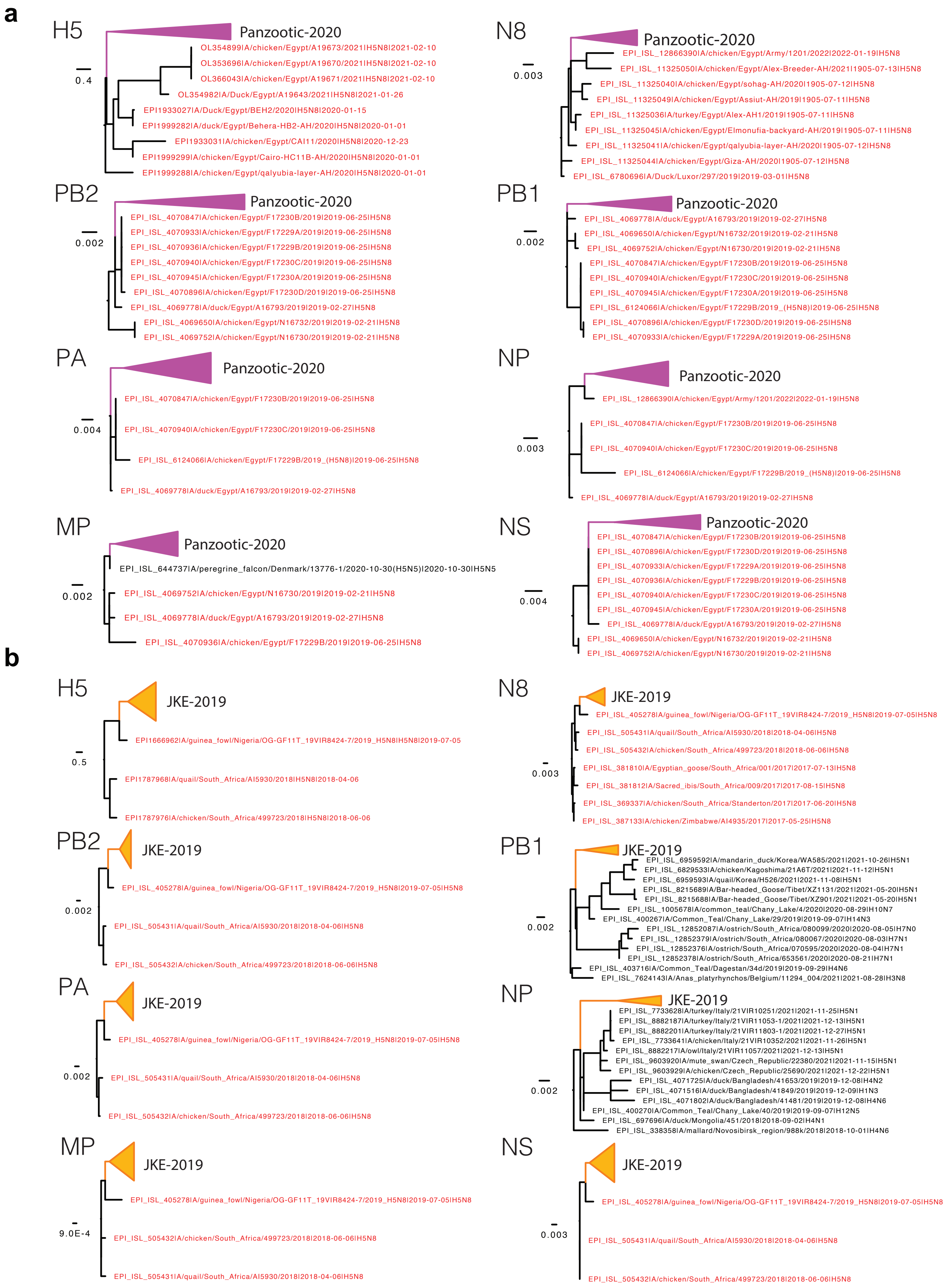


**Supplementary fig. 4.** Evolutionary relationships of Panzootic-2020 and JKE-2019 lineage. Maximum likelihood tree of Panzootic-2020 (a) and JKE-2019 (b) for eight segments. Samples collected in Africa are highlighted in red.
