## Supplementary Figure 5 for "The episodic resurgence of highly pathogenic avian influenza H5 virus"

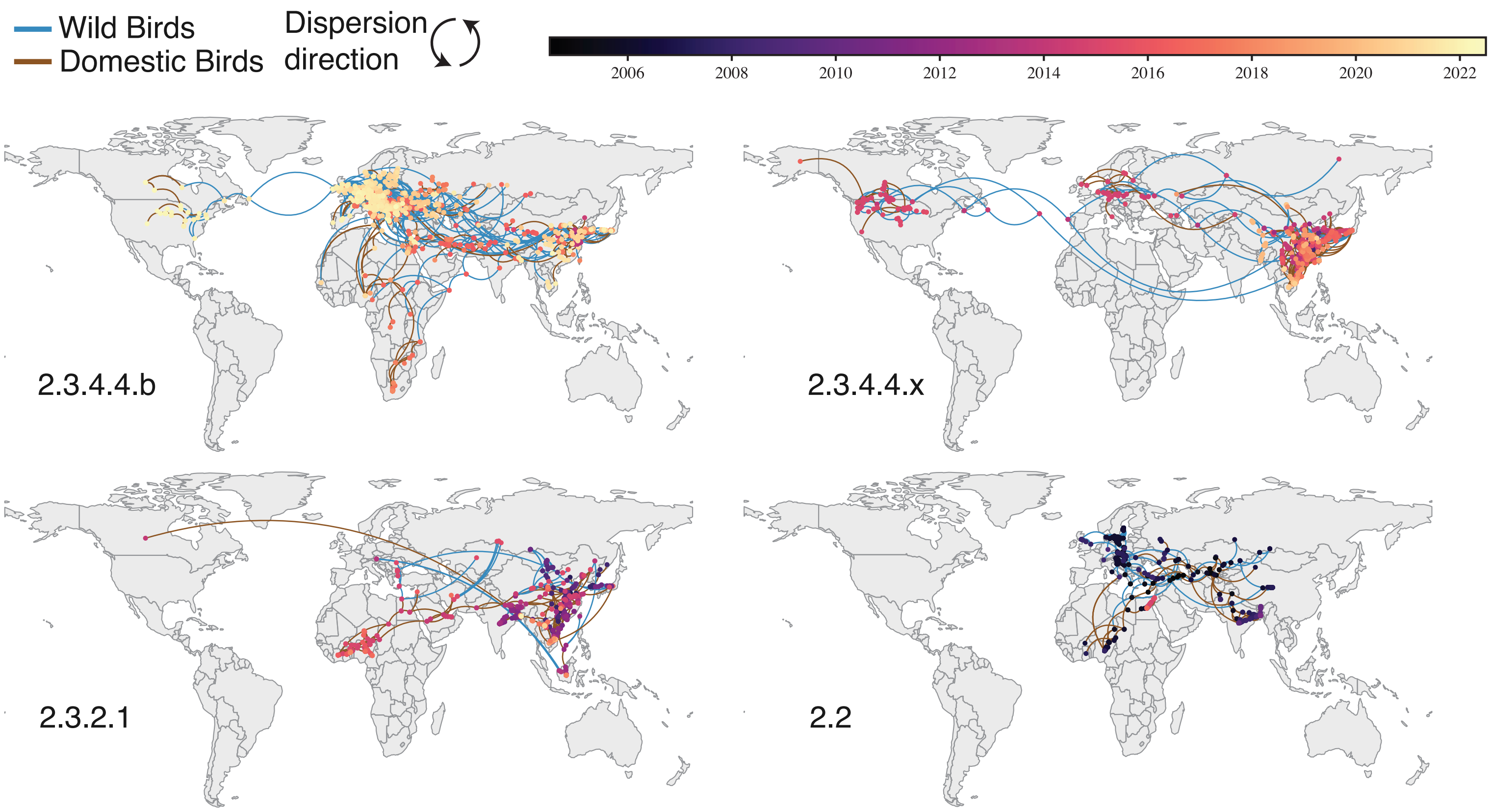


**Supplementary fig. 5.** Dynamics of HPAI H5 transmission lineages in clades 2.3.4.4.b, 2.3.4.4.x, 2.3.2.1 and 2.2. Virus lineage movements inferred by continuous phylogeographic analysis for each clade.
