## Supplementary Figure 6 for "The episodic resurgence of highly pathogenic avian influenza H5 virus"

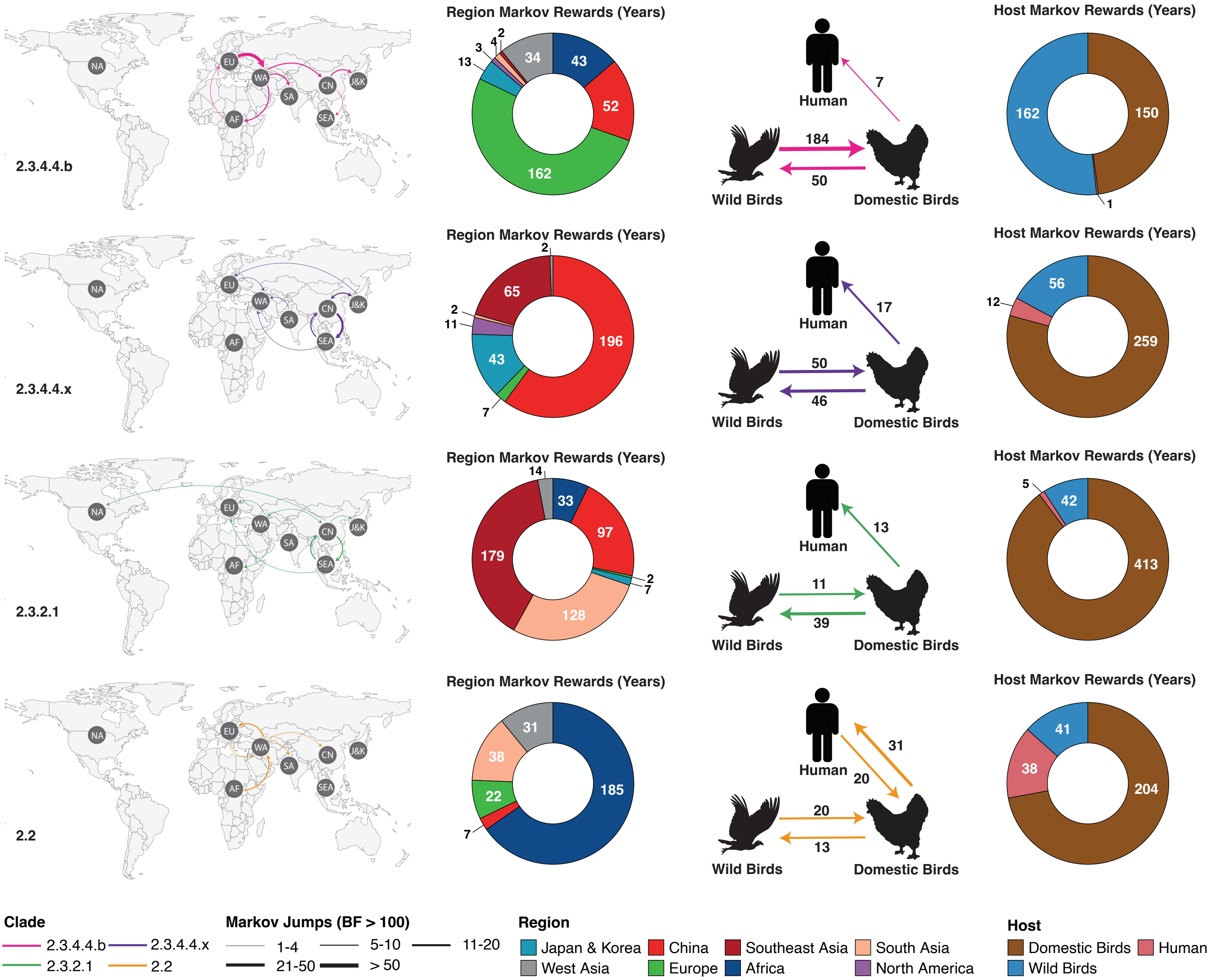


Supplementary fig. 6. The contrasting geographic and host transmission patterns among HPAI H5 2.3.4.4.b, 2.3.4.4.x, 2.3.2.1 and 2.2 clades inferred from discrete phylogeography. From left to right, the figures represent regional Markov jumps, regional Markov rewards, host Markov jumps and host Markov rewards.
