## Supplementary Figure 7 for "The episodic resurgence of highly pathogenic avian influenza H5 virus"

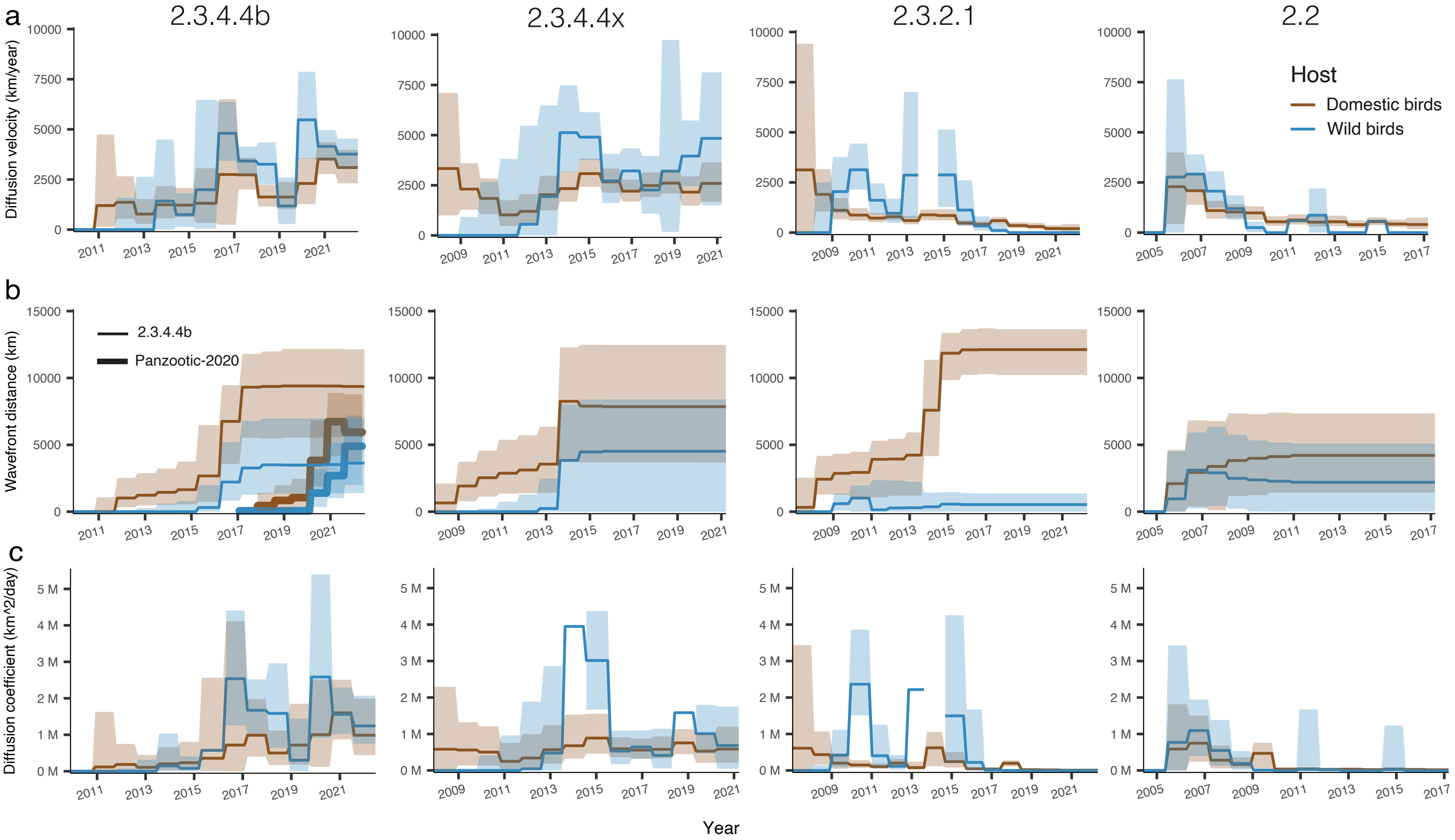


**Supplementary fig. 7.** The contrasting spatial epidemiology among HPAI H5 clades 2.3.4.4b, 2.3.4.4x, 2.3.2.1 and 2.2. (a) Viral Diffusion velocity (km/year) of domestic (brown) and wild birds (blue) over time for each clade. (b) Viral wavefront distance (km) of wild and domestic birds over time for each clade. A recalculation of the wavefront distance in the panzootic-2020 clade (including 2020/21 and 2021/22 resurgences, shown in a thick line) was performed, in which Egypt was regarded as the epidemic's origin. (c) Viral Diffusion coefficient (km$^2$/day) over time for each clade. The diffusion velocity and coefficient of wild birds in clade 2.3.2.1 during 2014 is not shown in the plot due to abnormal estimation value, possibly caused by insufficient sampling. The shaded areas denote the 95% confidence interval, in which some extreme values were not shown.
