## Supplementary Figure 8 for "The episodic resurgence of highly pathogenic avian influenza H5 virus"

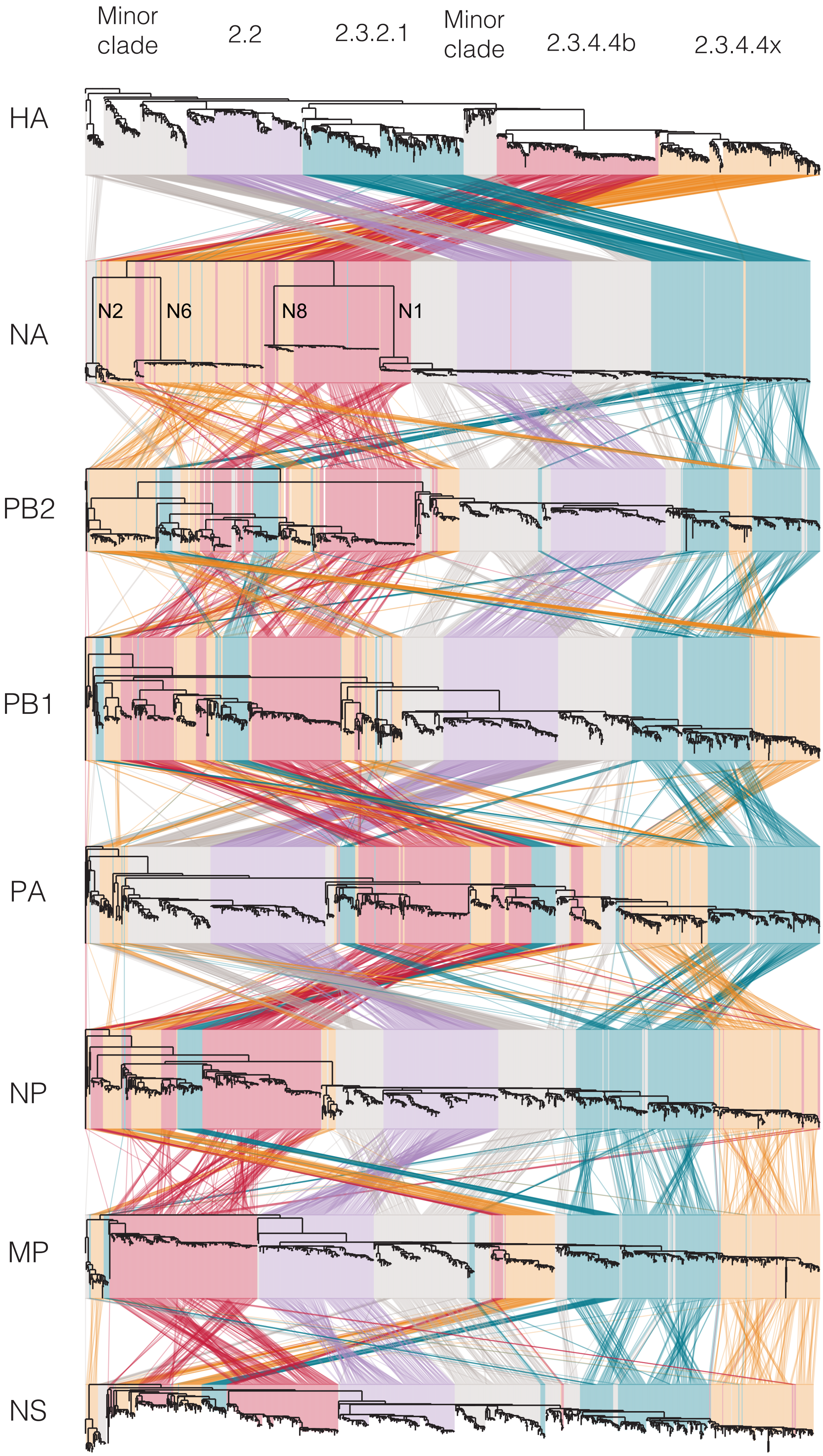


**Supplementary fig. 8.** Tanglegram of HPAI H5 virus reassortment. Coloured lines connect each virus across all eight gene, showing incongruence between and within major clades.
